# From GWAS Variant to Function: a Study of ~148,000 Variants for Blood Cell Traits

**DOI:** 10.1101/2021.02.16.431409

**Authors:** Quan Sun, Cheynna A. Crowley, Le Huang, Jia Wen, Jiawen Chen, Erik L. Bao, Paul L. Auer, Guillaume Lettre, Alexander P. Reiner, Vijay G. Sankaran, Laura M. Raffield, Yun Li

**Affiliations:** Department of Biostatistics, University of North Carolina at Chapel Hill, Chapel Hill, NC, USA; Curriculum in Bioinformatics and Computational Biology, University of North Carolina at Chapel Hill, Chapel Hill, NC, USA; Department of Genetics, University of North Carolina at Chapel Hill, Chapel Hill, NC, USA; Zilber School of Public Health, University of Wisconsin-Milwaukee, Milwaukee, WI, USA; Montreal Heart Institute, Montreal, Quebec, Canada; Department of Medicine, Faculty of Medicine, Université de Montréal, Montreal, Quebec, Canada; Department of Epidemiology, University of Washington, Seattle, WA, USA; Fred Hutchinson Cancer Research Center, University of Washington, Seattle, WA, USA; Division of Hematology/Oncology, Boston Children’s Hospital and Department of Pediatric Oncology, Dana-Farber Cancer Institute, Harvard Medical School, Boston, MA, USA; Broad Institute of Harvard and MIT, Cambridge, MA, USA; Department of Computer Science, University of North Carolina at Chapel Hill, Chapel Hill, NC, USA

**Keywords:** Genome-wide association studies, variant to function, functional annotations, experimental validations, blood cell traits

## Abstract

Genome-wide association studies (GWAS) have identified hundreds of thousands of genetic variants associated with complex diseases and traits. However, most variants are noncoding and not clearly linked to genes, making it challenging to interpret these GWAS signals. We present a systematic variant-to-function study, prioritizing the most likely functional elements of the genome for experimental follow-up, for >148,000 variants identified for hematological traits. Specifically, we developed VAMPIRE: Variant Annotation Method Pointing to Interesting Regulatory Effects, an interactive web application implemented in R Shiny (http://shiny.bios.unc.edu/vampire/). This tool efficiently integrates and displays information from multiple complementary sources, including epigenomic signatures from blood cell relevant tissues or cells, functional and conservation summary scores, variant impact on protein and gene expression, chromatin conformation information, as well as publicly available GWAS and phenome-wide association study (PheWAS) results. Leveraging data generated from independently performed functional validation experiments, we demonstrate that our prioritized variants, genes, or variant-gene links are significantly more likely to be experimentally validated. This study not only has important implications for systematic and efficient revelation of functional mechanisms underlying GWAS variants for hematological traits, but also provides a prototype that can be adapted to many other complex traits, paving the path for efficient variant to function (V2F) analyses.

## Introduction

Genome-wide association studies (GWAS) have identified thousands of genetic loci and hundreds of thousands of genetic variants associated with various complex human diseases and traits, but the underlying genetic mechanism for the vast majority of these GWAS signals remains elusive. With extensive sequencing and GWAS efforts, there is a pressing need to convert the large and ever growing number of significant GWAS variant-trait pairs into human-interpretable functional or mechanistic knowledge^1^. Most variants identified through GWAS reside in the noncoding regions (e.g., >95% for blood cell traits^2^), and most signals include multiple highly correlated variants or variants in strong linkage disequilibrium (LD). Pinpointing the most likely causal variants within GWAS signals, and linking these variants to their target genes, is challenging, particularly as the number of GWAS loci and variants increases. For hematological traits, for instance, our recent GWAS meta-analyses^3;4^ have revealed over seven thousand loci, with >148,000 variants associated with at least one blood cell index at stringent genome-wide significance threshold. Comprehensive and computationally efficient annotation and prioritization of such GWAS findings are of ever-increasing interest.

Understanding how genetic variants contribute to a phenotype is often referred to as the variant-to-function (V2F) problem. Responding to this problem requires us to determine causal genetic variants, relative cell types/states, their target genes and cellular/physiological functions^5^. Functional experiments are needed to fully reveal molecular mechanisms, but we cannot yet afford to perform time-, money- and labor-consuming experimental validations of thousands of loci involving hundreds of thousands of potentially functional variants or regulatory elements controlling their nearby genes, since each gene is likely regulated by multiple variants and each variant may regulate multiple genes. Thus, computational methods are needed to screen potential variants and their effector genes for further experiments.

In this study, we focus on hematological traits. Hematological phenotypes (red blood cell, white blood cell, and platelet counts and indices) are critical physiological intermediaries in oxygen transport, immunity, infection, thrombosis, and hemostasis and are associated with autoimmune, allergic, infectious, and cardiovascular diseases. Hematological traits are highly heritable ^6^, and recent large GWAS for hematological traits (including nearly 750,000 participants) identified thousands of variant-trait associations ^2;4^ In addition, there are multiple large-scale functional experiments already available^2;7;8^ for hematological traits, as well as fairly comprehensive functional annotation resources relevant to blood tissues. This makes hematological traits an ideal model for this type of V2F computational solution.

We have developed VAMPIRE: Variant Annotation Method Pointing to Interesting Regulatory Effects, a tool for the user to explore annotations encompassing epigenomic signatures, variant impact on protein and gene expression, chromatin conformation information from Hi-C and similar technologies, as well as publicly available GWAS and PheWAS results, creating a comprehensive annotation profile for variants from recent trans-ethnic blood cell trait publications^3;4^ with a flexible interface for adding additional future GWAS results. This interactive web application implemented in R Shiny provides a model display mechanism for annotating GWAS variants from diverse complex traits, allowing selection of most likely causal variants and their effector genes for experimental follow-up. Importantly, we show the value of how variants and genes nominated by VAMPIRE can highlight key regulators of blood cell traits using independent functional assessment, confirming the value of this annotation tool. While blood cell traits are the focus for VAMPIRE, this framework (including our R Shiny application) is adaptable for annotation of other complex trait GWAS results and will facilitate the connection between variant and function.

## Methods

### Variant Annotations

The current version of VAMPIRE includes GWAS results from two studies (as detailed in Supplemental Methods), including all variants in 95% credible sets for fine-mapped hematological trait associated loci from Chen et al. (N=148,019 variants) ^4^ and lead variants (N=2) from a TOPMed imputed GWAS meta-analysis in African American and Hispanic/Latino populations^3^. We plan to extend VAMPIRE as new trans-ethnic blood cell trait genetic analyses are released.

The sources of the annotation used are stated clearly in the VAMPIRE online application, with links or references to the original data sources. As a brief summary, the annotation categories are trivially split into six types (“variant level”, “1D”,“2D”,”3D”, “PheWAS”,“GWAS”). First, “variant level” contains data on phenotypic association from the original publication or preprint (such as the p-value for association with a given hematological trait, effect size, and posterior probability of inclusion for fine-mapping credible sets). Second, “1D” refers to epigenomic or sequence constraints features. This displays selected output from WGSA ^9^ including functional prediction scores, conservation scores, and epigenetic information gathered from GeneHancer ^10^, FANTOM5 ^11;12^, Roadmap ^13^, and ENCODE ^14^ ATAC-seq peaks from recent studies for blood cell traits ^15;16^ and key histone ChIP-seq peaks such as H3K9me3, H3K36me3, H3K4me1, H3K4me3, and H3K27Ac generated across blood cell related tissues from Roadmap Epigenomics are also included ^13;17^ We further include information regarding whether each variant resides in any selective sweep region detected from multiple populations in the 1000 Genomes Project^18^ using the S/HIC method^19;20^. Information is displayed based on the tissue relevance to the blood cell phenotype (see Supplemental Methods). All variants have 1D annotation, but for prioritization purposes as described below in the five categories for noncoding variant annotation, we define 1D annotation as FANTOM5_enhancer_robust =Y (yes), or Genehancer_feature=“Promoter” or “Enhancer” or “Promoter/Enhancer”, or coreMarks (for any relevant roadmap epigenomic category) =“Enhancers” or “Active TSS.” Users can then additionally filter by criteria such as functional prediction and conservation scores.

For the “2D” annotations, we included impact on gene expression and splicing ratios (eQTL and sQTL information) and impact on protein abundance (pQTL information^21^) from public sources relevant to blood cell traits. This includes both bulk and cell type specific sources from the public domain (eQTLGen ^22^, CAGE ^23^, BIOS ^24^ for whole blood, and Raj et al for purified CD4+ T cells and monocytes ^25^). Information available in these sources varies, but generally we at a minimum display the effect size estimate, p-value, the allele assessed, and the gene or protein involved. Variants were matched across sources based on chromosome, position, and alleles of each variant. Only significant results (based on FDR or other publication specific thresholds) from the respective sources are displayed in VAMPIRE; we do note that formal co-localization analyses would still need to be performed to determine if blood cell related and gene/protein expression QTL signals truly coincide.

For the “3D” annotations, we include information on 3D genome conformation, linking blood lineage specific regulatory elements to target genes from various sources. More specifically, using Hi-C data we incorporated statistically significant long-range chromatin interactions (LRCI) ^17;26;27^ calculated from Fit-Hi-C ^28^, loops using the HiCCUPs methodology ^26^, and super-FIREs for related tissues^17^. Two Promoter-Capture Hi-C (PCHi-C) data sources^29;30^ were also incorporated and matched with the 2D results to highlight consistent evidence regarding the affected gene(s) across “2D” and “3D” annotations. VAMPIRE displays information on the number of loops, LRCI, PCHi-C interactions, FIREs, or super-FIREs, as well as significance measures such as p-values, FDR, or CHICAGO scores where applicable. This “3D” annotation information can also be visualized via our HUGIn browser ^31^.

The last two data groups present results from two PheWAS sources ^4;32^ and GWAS results of blood cell traits from GWAS catalog ^33^, allowing the user to evaluate if hematological trait associated variants may also influence other complex traits.

To visualize and leverage these multiple annotation categories for further analysis or prioritization of experimental validations, VAMPIRE efficiently displays and integrates relevant variant information, allowing the user to investigate either all the variants annotated or subsets based on annotation category groupings, searching either by variant or by gene name. The comprehensive annotation for the variants is summarized using a five category grouping created for highlighting the most promising variants as they have various types of annotation. Specifically, the five categories for noncoding variants are (1) the most restrictive category, containing variants that have 1D, 2D, and 3D annotation and the genes implicated by 2D and 3D evidence are consistent; (2) containing variants with 1D, 2D, and 3D evidence, but the genes implicated from different resources are not consistent; (3) 2D and 3D with consistent gene evidence between the 2D and 3D annotations; (4) variants with 2D and 3D information and no consistent gene implied; (5) variants with 1D and 3D evidence. We also have a predicted high impact coding variant category displayed, including high confidence loss of function (LoF) variants and likely influential missense, in frame indels, and synonymous variants. Variants without strongly compelling variant annotation are still displayed, but are not listed in these high priority categories. The user can further subset results by hematological trait, hematological trait category, or (for the Chen et al paper ^4^) the ancestry specific grouping in which a given credible set was derived (trans-ethnic, European, East Asian, South Asian, Hispanic/Latino, or African ancestry). In addition, the user can restrict the amount of information presented by selecting which tables to be displayed. All tables can be exported in a csv or tab delimited format.

### Enrichment analysis

To assess whether the variants prioritized by VAMPIRE are more likely to be functionally impactful, we performed enrichment analysis at three different levels: variant level, gene level, and variant-gene pair level, leveraging data generated from previously published functional experiments ^2;7;8^. For each set of analysis, we conducted Fisher’s exact test and calculated odds ratios (OR) and one-sided p-values.

At the variant level, we assessed the enrichment of variants that modify transcription factor (TF) binding motif^2^ among our annotation category 1 variants. Recently, Vuckovic et al. ^2^ characterized variants that affect erythropoiesis or hematopoiesis by modifying related TF motifs, such as for KLF1, KLF6, MAFB, and GATA1. We chose these four erythroid TFs as positive control TFs and two non-erythroid TFs (IRF1 and IRF8) as negative controls.

At the gene level, we evaluated the genes interrogated by Nandakumar et al. ^8^ with a pooled short hairpin RNA (shRNA) based loss-of-function approach. Specifically, Nandakumar et al. studied 389 candidate genes in the neighborhood of 75 loci associated with red blood cell traits ^34^, to identify potential causal genes underlying these GWAS signals. We assessed the enrichment of genes validated by shRNA experiments among those prioritized in VAMPIRE’s category 1. Note that the categories were previously defined at variant level. Here we extent variant category to gene category as the strongest category where a genome-wide significant variant linked to this gene falls in.

At the variant-gene pair level, we employed the enhancer-gene connections validated via CRISPRi-FlowFISH experiments by Fulco et al. ^7^ in their activity-by-contact (ABC) paper. Specifically, Fulco et al. tested pairs of candidate *cis* regulatory elements (CREs, ~ 500bp regions) and their potential effector genes via CRISPRi perturbations of the CREs, in multiple cell lines including the K562 cells. Fulco et al. tested 4,124 CRE-gene pairs in total, of which 175 were significant from their experiments. We overlapped their tested CREs with variants in our VAMPIRE annotation database. We define a VAMPIRE variant-gene pair confirmed if the variant overlaps an ABC validated CRE *<u>and</u>* the linked genes in VAMPIRE (from QTL and chromatin capture conformation evidence) overlaps the corresponding effector gene for that CRE via ABC’s CRISPRi-FlowFISH experiment. We focused on ABC experiments performed on the K562 cells (instead of GM12878 cells, where a very small number of CREs were tested) as the number of tested CRE-gene pairs was not too small for robust statistical inference. Matching the K562 cell line, we focused only on variants associated with red blood cell traits. Similar to the above two sets of enrichment analyses, we focused on annotations in VAMPIRE’s prioritization category 1. Specifically, we tested whether variant-gene pairs prioritized in VAMPIRE’s category 1 are enriched within ABC’s validated enhancer-gene connections. Given the CREs tested in the ABC paper are rather short (~ 500bp), we also performed sensitivity analysis by first extending the CRE regions by +/− 1kb and +/− 5kb and then overlapping variants with these extended CREs, to ensure robust conclusions.

## Results

### Overview of VAMPIRE annotations

The overall framework of VAMPIRE is illustrated in **Figure 1**. We started with all variants in 95% credible sets from our recent trans-ethnic study for hematological traits (total 148,019 variants) ^4^ and lead variants (2 variants) from Kowalski et al.^3^. We incorporated six types of annotations (detailed in **Methods**): GWAS summary statistics and posterior probability of inclusion from our previous fine-mapping analyses ^4^; epigenomic or sequence constraints features (1D); eQTL, sQTL and pQTL information (2D); information on 3D genome conformation (3D); results from two PheWAS sources^4;32^ (PheWAS); and GWAS results from blood cell traits from GWAS catalog ^33^ (GWAS).

**Figure 1.**
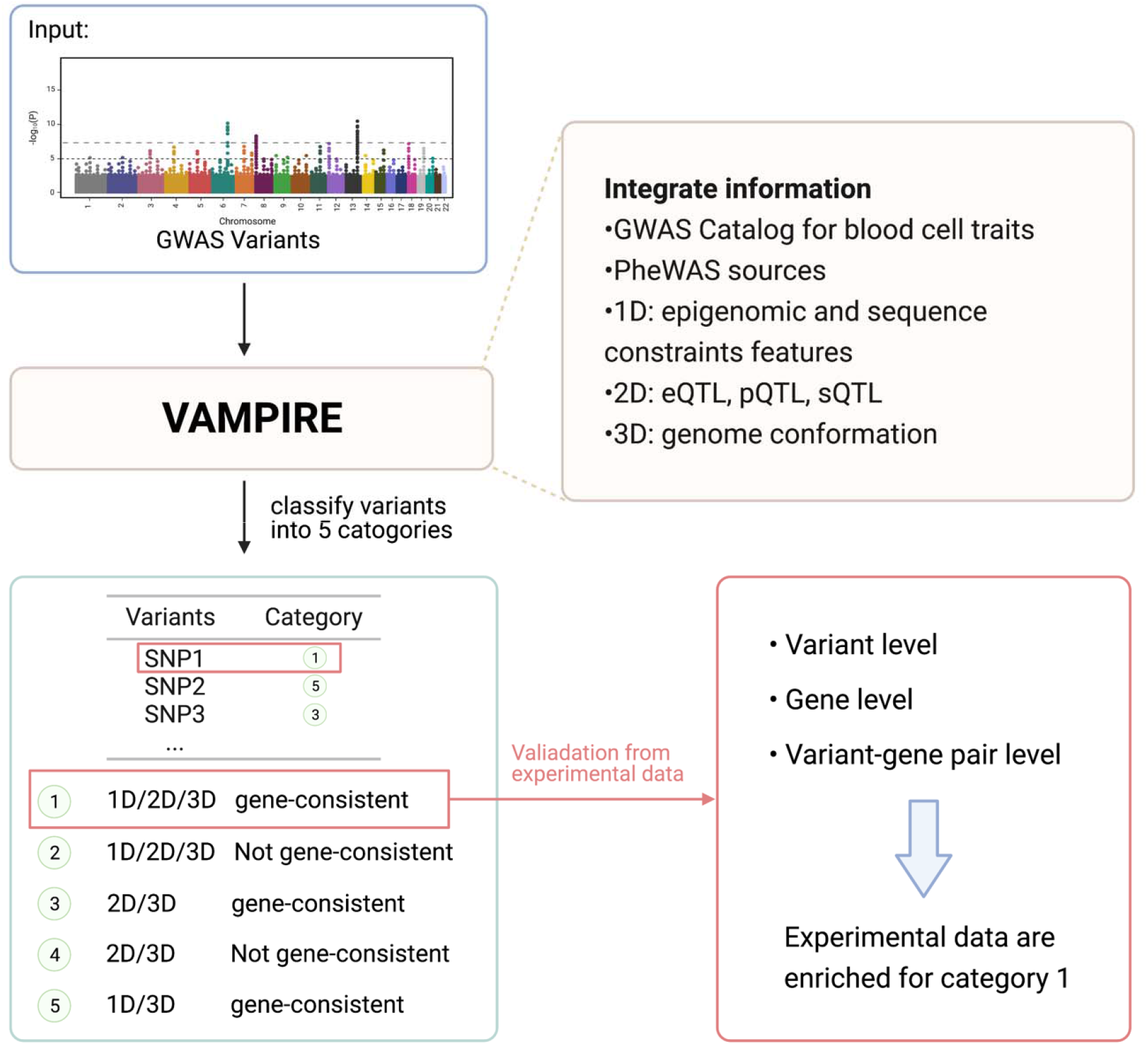
Overall framework of this study. VAMPIRE starts with GWAS variants in the 95% credible sets, integrates different annotations and assigns them into different prioritization categories. We further demonstrated that our top prioritized category is enriched with variants that were experimentally validated. VAMPIRE provides a prototype that can be adapted to many other complex traits, paving the path for efficient variant to function (V2F) analyses.

To visualize and prioritize variants, their corresponding candidate regulatory regions, and their potential effector genes, we leverage the aforementioned six types of annotation to group these ~ 148,000 variants into various prioritization categories. Specifically, for non-coding variants, we classified them into five categories (detailed in Methods). Among them, category 1 is the most restrictive category, containing variants that have 1D, 2D, and 3D annotation and the genes implicated by 2D and 3D evidence are consistent. Variants not falling into any of the five categories are classified as uncategorized. In addition, each gene is categorized according to the prioritization categories of its linked variant(s). When its linked variants fall in multiple categories, the gene is assigned to the most highly prioritized category.

### Enrichment analysis

Our enrichment analyses employing multiple previously published functional validation experiments encompassing variant-level, gene-level, and variant-gene pair levels all showed promising results. Specifically, at the variant level, we found significant enrichment of variants affecting TF binding motifs among variants prioritized in category 1 of VAMPIRE (**Figure 2**), for all the erythroid TFs (p < 8.1E-4) but GATA1 (p = 0.18) (**Table 1**), likely due a smaller sample size of variants. In contrast, neither of the two negative control TFs (IRF1 and IRF8) showed any significant enrichment (p = 0.22 and 0.62). At the gene level, we focused on two statistics: (1) number of genes selected for shRNA experiments, since genes were more likely to be selected for experiments when they demonstrated some prior evidence of potential causality, and (2) number of genes validated (p < 0.05) by shRNA experiments. We compared the number of genes in our annotation category 1 and all other categories, and found that both shRNA candidate genes (p = 3.5E-13) and significant genes (p = 3.1E-8) show strong enrichment among those in our annotation category 1 (**Table 2**), and the estimated enrichment score for significant genes (OR = 4.65) is almost double of that for candidate genes (OR = 2.37). These results suggest the genes prioritized by VAMPIRE’s category 1 annotations are more likely to be functional.

**Figure 2.**
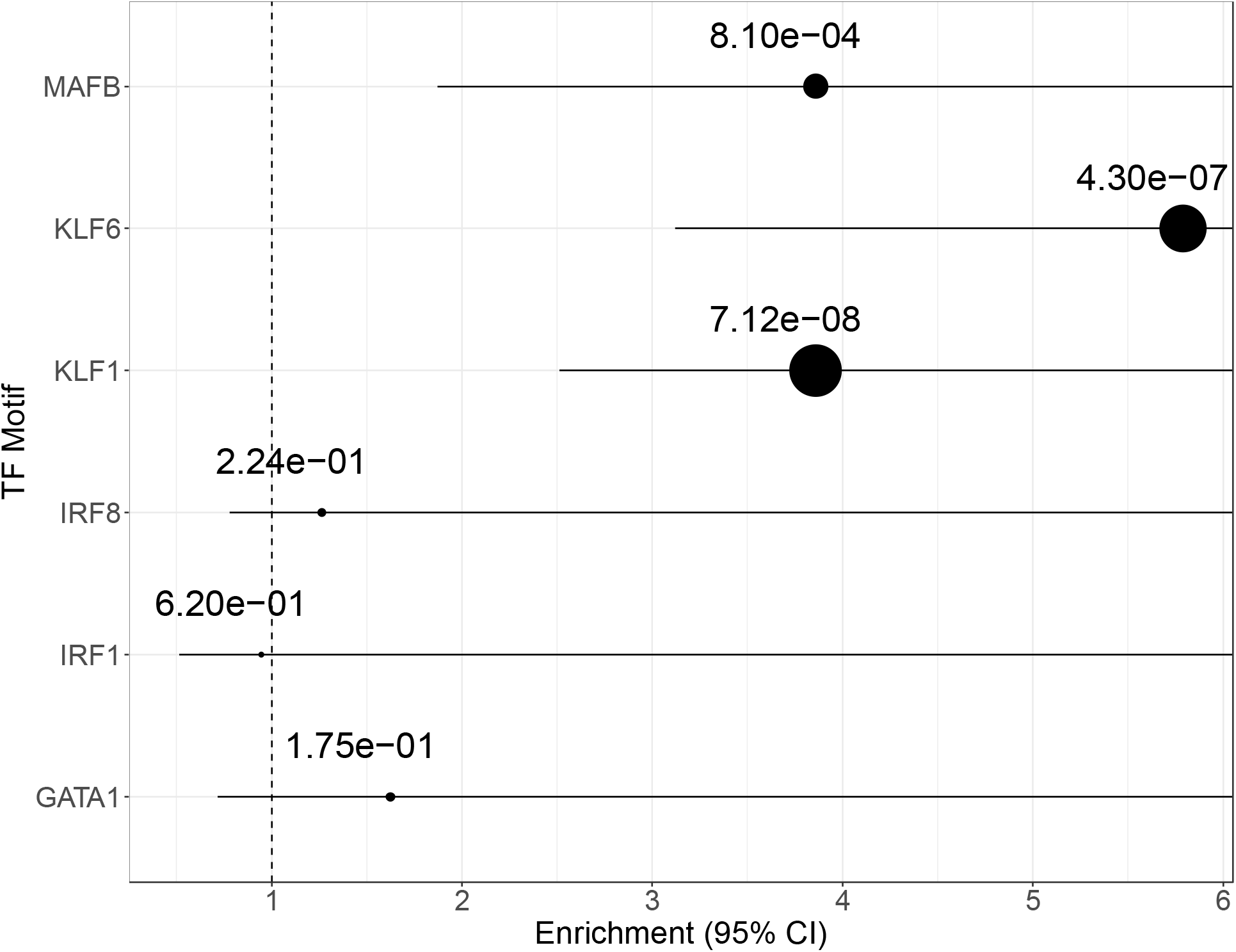
Variant level TF motif enrichment analysis. Each dot represents an enrichment score with the line depicting 95% confidence interval (CI). All the upper bounds of these CIs are infinity. The p-values of the enrichment are reflected by the dot size at the OR point estimate with a larger dot indicating more significant the enrichment.

**Table 1.**
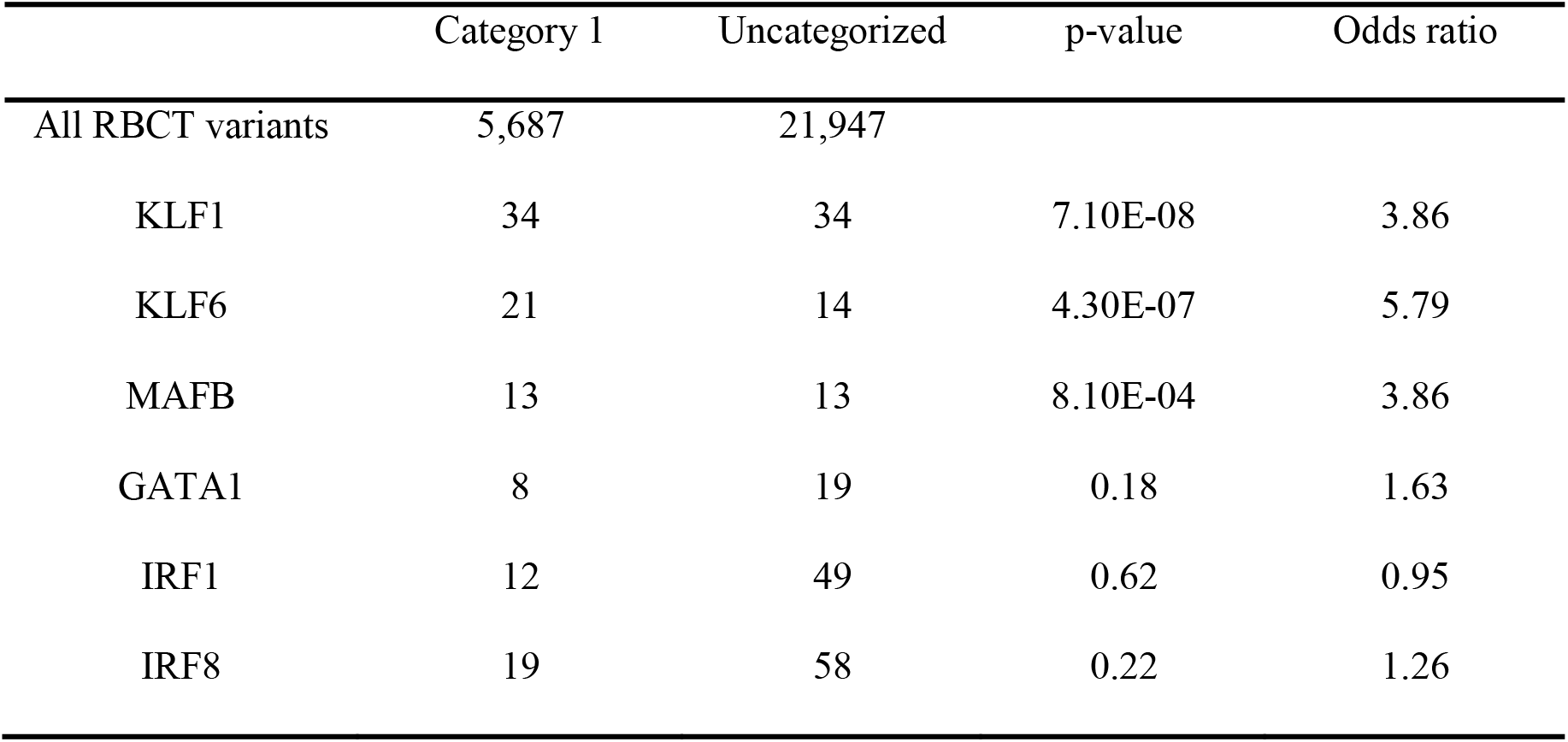
Variant level transcription factor (TF) motif enrichment analysis. Four erythroid TFs and two non-erythroid TFs were selected. Fisher’s exact test was applied to test for enrichment. Three erythroid TFs show enrichment for our VAMPIRE annotation category 1 (MAFB, KLF6, KLF1, p<0.05). GATA1 motif variants also have some evidence of enrichment (odds ratio = 1.625) but this enrichment is not significant (p=0.18), likely due to smaller sample size of variants. Two non-hematopoiesis transcription factors selected as controls don’t show significant enrichment with VAMPIRE functional annotation category 1. RBCT, red blood cell trait associated.

|  | Category 1 | Uncategorized | p-value | Odds ratio |
| --- | --- | --- | --- | --- |
| All RBCT variants | 5,687 | 21,947 |  |  |
| KLF1 | 34 | 34 | 7.10E-08 | 3.86 |
| KLF6 | 21 | 14 | 4.30E-07 | 5.79 |
| MAFB | 13 | 13 | 8.10E-04 | 3.86 |
| GATA1 | 8 | 19 | 0.18 | 1.63 |
| IRF1 | 12 | 49 | 0.62 | 0.95 |
| IRF8 | 19 | 58 | 0.22 | 1.26 |

**Table 2.** Gene level enrichment analysis. Fisher’s exact test was applied to test for enrichment. Both shRNA experiment candidate genes and validated genes show significant enrichment in our most restrictive VAMPIRE annotation category (category 1).

|  | Category 1 | Other categories | p-value | Odds ratio |
| --- | --- | --- | --- | --- |
| All category genes | 9,857 | 7,408 |  |  |
| shRNA Candidate genes | 262 | 83 | 3.50E-13 | 2.37 |
| shRNA Validated genes | 68 | 11 | 3.10E-08 | 4.65 |

Finally, at the variant-gene pair level, we also observed enrichment among variants selected into VAMPIRE’s category 1 (**Table 3**). When restricting only to variants in category 1 and associated with red blood cell traits and without extending the CRE regions, only 7 of VAMPIRE’s variant-gene pairs can be found in ABC’s CRISPRi-FlowFISH experiments, of which 6 are not significant and 1 is significant. While not significant (p = 0.26), the direction of enrichment is nevertheless encouraging (one of seven, or 14.3%, confirmed by CRISPRi-FlowFISH experiments) and 3-fold greater than that among all/background pairs from Fulco et al. ^7^, where 175 out of 4124 variant-gene pairs (4.2%) were confirmed. Note that all the confirmed pairs were linked with variants associated with red blood cell traits. Further generalizing to all VAMPIRE annotation categories and to variants associated with any blood cell trait, the enrichment OR increases to 8.30 with p-value 9.0E-5, indicating that variant-gene pairs prioritized by VAMPIRE’s five categories have much higher odds of being functional. To further accommodate causal variants tagged by GWAS variants not falling into the short 500bp CREs, we extended the CREs by +/− 1kb or +/− 5kb, and performed similar enrichment analysis. Our conclusions remained qualitatively similar (**Table 3**), but the enrichments increased in significance, thanks to larger sample size (in this context, the larger number of variant-gene pairs contributing to the analysis) and suggesting that more liberal windows of *cis*-regulatory regions can capture a higher rate of functional variant-gene pairs. For example, the enrichment for category 1 variants associated with red blood cell (RBC) traits reached an OR of 15.77 (p=3.8E-6) and 16.68 (p=3.1E-15) for 1kb and 5kb extension, respectively. We thus conclude that such enrichment is significant and robust to the extension of CREs.

**Table 3.**
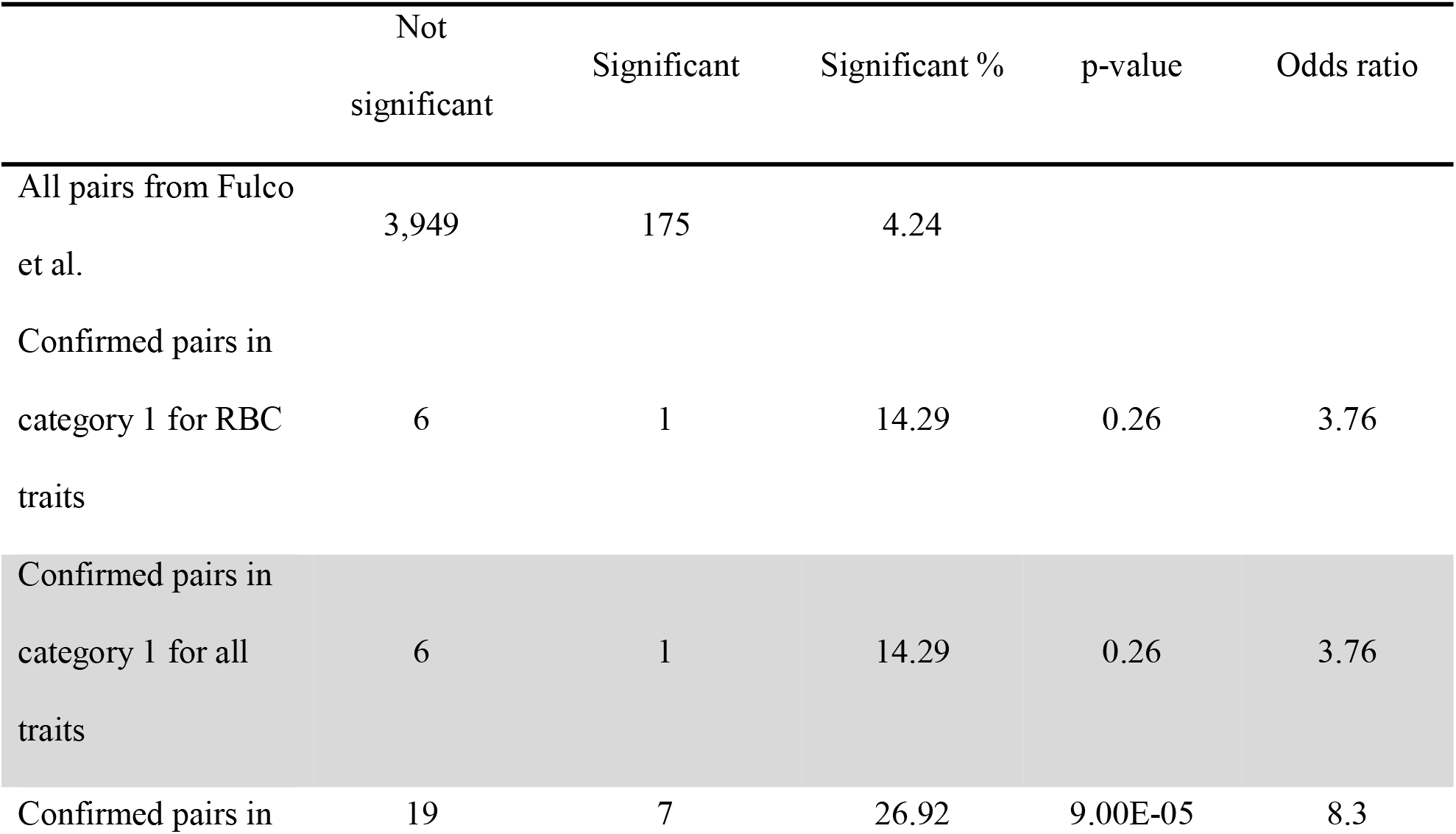
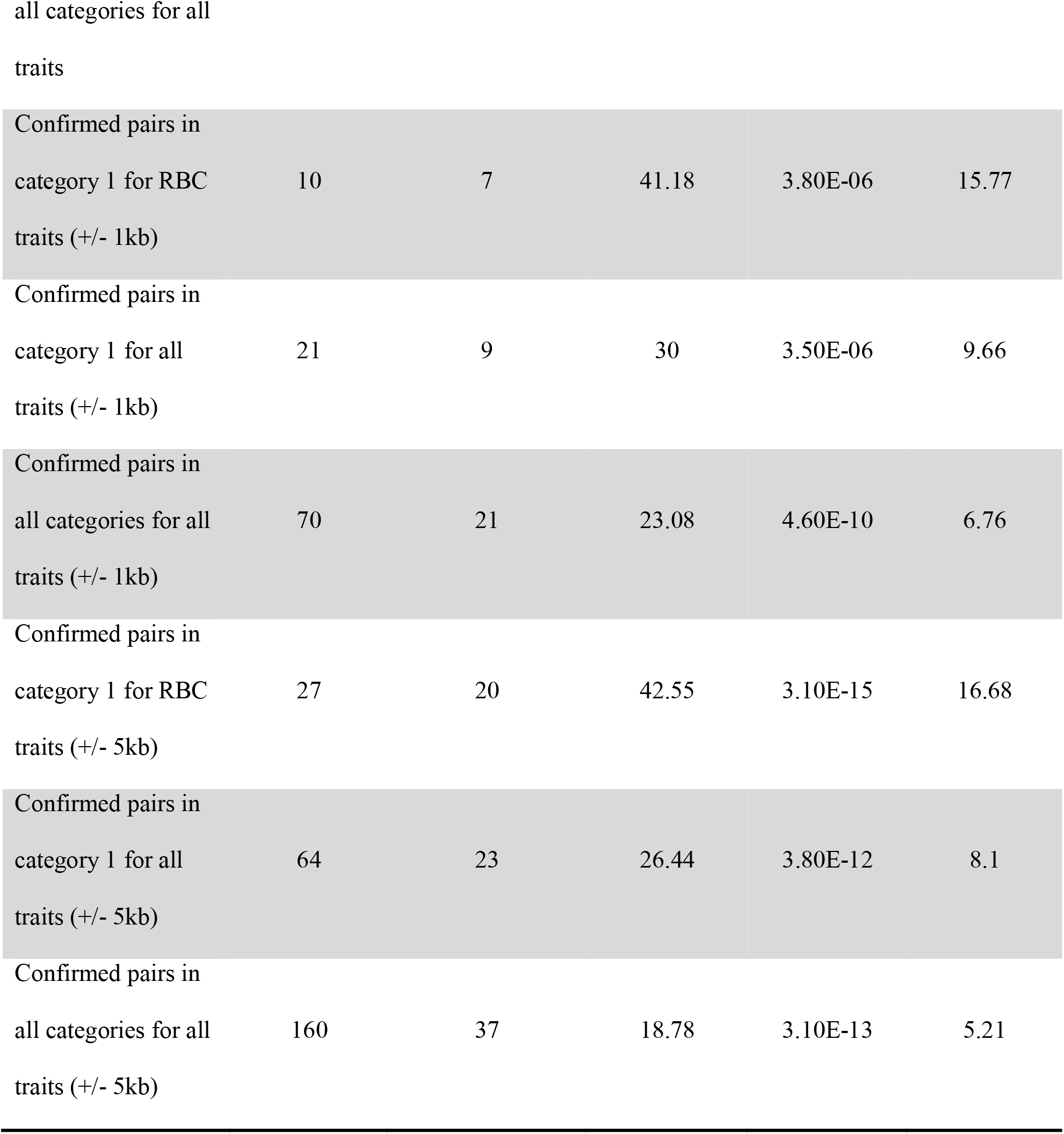
Variant-Gene pair level enrichment analysis. We performed analysis for three variant annotation pools (category 1, red blood cell (RBC) trait associated; category 1, any blood cell trait associated; any annotation priority category (1-5), any blood cell trait associated) and three CRE lengths. Fisher’s exact test was applied to test for enrichment. We found enrichment for all three variant annotation pools. These enrichments are also robust to the extension of CREs.

### Application example

**Figure 3** shows one example at the *CALR* locus associated with red blood cell traits. Fulco et al. confirmed by CRISPRi-FlowFISH experiment that CRE chr19:12,996,905-12,998,745 (hg19) regulates gene *CALR* (adjusted p-value 1.9E-7)^7^. Annotations compiled by VAMPIRE suggest, consistently, that *CALR* is linked to rs8110787 (chr19:12,999,458, hg19) in category 1. rs8110787 is associated with several RBC traits ^4^, including hematocrit (HCT), mean corpuscular hemoglobin (MCH), mean corpuscular volume (MCV) and red blood cell counts (RBC). Based on genomic distance alone, *CALR* is not the nearest gene to rs8110787, with several other closer genes. However, based on H3K27ac HiChIP data in K562 cells ^35^, rs8110787 significantly interacts with *CALR* promoter region (p < 1E-120), suggesting that *CALR* is a potential target gene regulated by the CRE around rs8110787. This variant is also an eQTL of *CALR* from CAGE ^23^ (p = 9.4E-16) and BIOS ^24^ (p = 1.0E-25), and is an enhancer in K562 Leukemia cells (E123) from Roadmap ^13^, adding additional evidence. Our VAMPIRE successfully highlights this rs8110787-*CALR* pair in its category 1.

**Figure 3.**
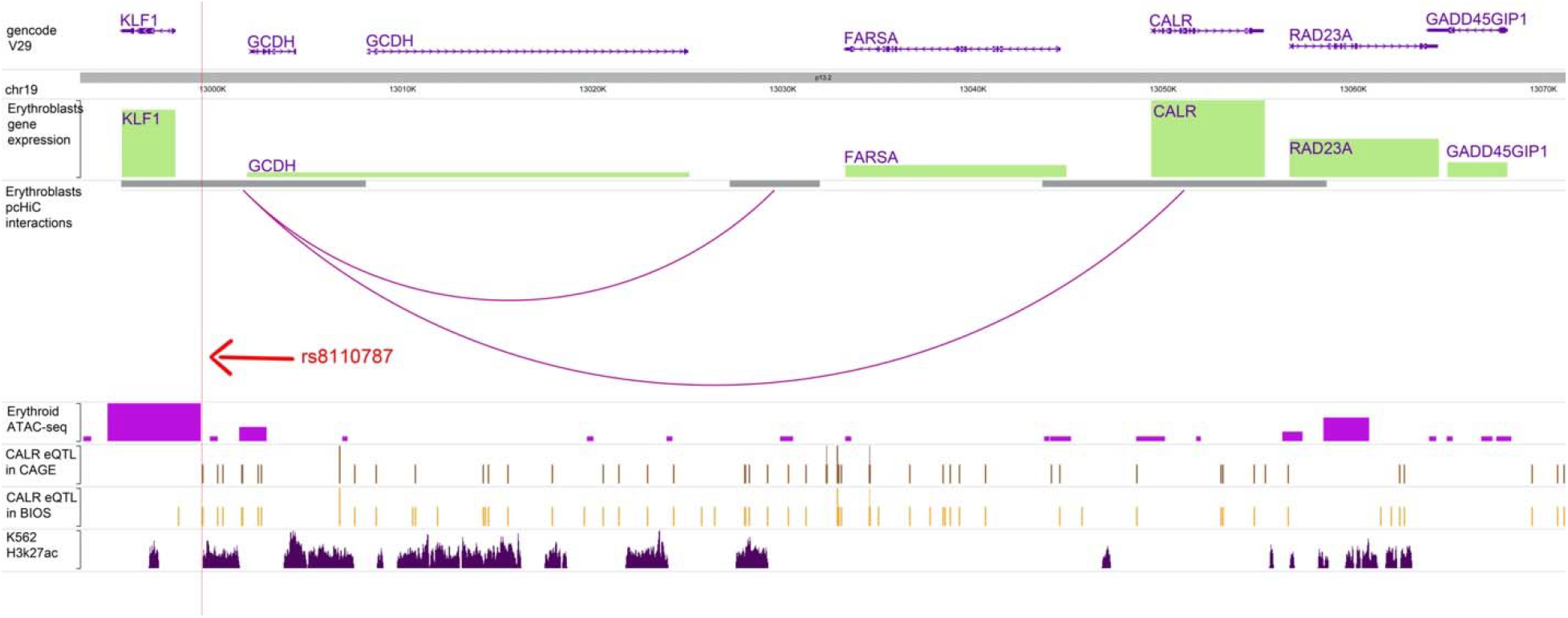
Variant-gene pair example (*rs8110787-CALR*) visualization from HUGIn2 ^31^. Fulco et al. confirmed via CRISPRi experiments that chr19:12996905-12998745 (hg19) regulates gene *CALR* (adjusted p-value 1.9E-7) which is highly expressed in Erythroblasts^7^. Based on annotations in VAMPIRE, *CALR* is linked to rs8110787 (chr19:12999458, hg19) in prioritization category 1, including higher than expected physical interactions with the *CALR* locus from erythroblasts pcHiC data^29^, eQTL of *CALR* in CAGE^23^ and BIOS^24^, erythroid ATAC-seq peak^16^ and H3K27ac peak in K562 leukemia cell^13^. rs8110787 is associated with several RBC traits (namely hematocrit, mean corpuscular hemoglobin, mean corpuscular volume, and red blood cell count) as reported in Chen et al. ^4^.

As a further example of the utility of the VAMPIRE application, we present the annotation results for one of the lead genome-wide significant variants from recent trans-ethnic GWAS analyses from Chen et al. ^4^ For our analysis, we were particularly interested in exploring low frequency variants, and those more common in those of non-European ancestry. We were able to quickly rank and prioritize variants for further examination using the annotation categories described above, including the low frequency variant rs112097551 associated with mean corpuscular volume (MCV), mean corpuscular hemoglobin (MCH), and red blood cell count.

This low frequency intergenic variant rs112097551 (*GATA2-RPN1* locus, 0.15% minor allele frequency in Chen et al. trans-ethnic analysis ^4^) has no close linkage disequilibrium proxies in African or European populations, and thus was not compared to other highly correlated variants. Based on variant frequency, particularly in European ancestry populations, we had no expectation this variant would have eQTL or pQTL evidence (2D annotation), given currently available sample sizes for eQTL and pQTL analysis. For low frequency variants, 1D and 3D annotation would be the highest annotation category likely for a variant of interest like rs112097551. The variant is ~ 5x more common among African versus non-African samples in gnomAD version 2.1.1. It is the only variant in the credible set in fine-mapping analyses from Chen et al. 1D annotation suggests this variant is highly conserved (CADD Phred score of 20.4, meaning the variant is amongst the top 1% of deleterious variants in the human genome), and it is rated as deleterious by FATHMM-XF (rank score 0.99169, close to the maximum score of 1). It is also in open chromatin in megakaryocyte–erythroid progenitor cells, based on hematopoietic ATAC-seq data ^36^. 3D annotation from PCHi-C data in erythroblasts from Javierre et al. ^29^ links this variant to the gene *RUVBL1* ~ 500Kb away, as well as noncoding transcripts *RNU2-37P* and *RUVBL1-AS1*. Based on this data, which can be quickly displayed using the VAMPIRE application, we are currently working on *in vitro* follow-up of this candidate functional enhancer variant^37^.

## Discussion

As genotyped sample sizes increase and meta-analysis efforts grow ever larger, more variant-trait pairs are identified for complex traits than can be easily annotated on a variant by variant basis. New, user-friendly applications are needed for rapid display of functional annotation information and prioritization of variants for further functional follow-up to pave the V2F path. Our VAMPIRE tool provides an example of how the publicly available code can be adapted to accommodate other sources of annotation specific to other complex trait GWAS results or to accommodate future blood cell trait GWAS and annotation resources. In addition to *a priori* providing one category of coding variants and 5 categories of non-coding variants that warrant prioritization consideration, VAMPIRE allows users to decide their own categories based on arbitrary combinations of the annotations at adjustable thresholds (for example, prioritizing high CADD score variants, or variants in open chromatin in blood cells based on ATAC-seq). Along with the addition of more blood cell trait genetics papers published in the future, VAMPIRE could also be used as written to annotate GWAS results for other blood related phenotypes, such as recent GWAS of risk of myeloproliferative neoplasm or clonal hematopoiesis ^38;39^.

As we accumulate additional functional validation data, including high-throughput massively parallel reporter assays (MPRA), medium-throughput CRISPRi/CRISPRa and low throughput mouse xenotransplant experiments, we will provide statistics summarizing experimental validation results (e.g., number of variants in the category followed-up, proportion that show evidence of functional impact in their experiments) for each of the 6 VAMPIRE categories and for user defined categories. Importantly, we illustrate the value of VAMPIRE using existing independent functional validation and therefore illuminate the value of this type of annotation tool in enabling one to go from variant to function for blood cell traits and other complex phenotypes.

We also note that there are some limitations of VAMPIRE. First, comprehensive annotations specific to various cell types and cell states would further enhance classification and prioritization accuracy of functional variants or regulatory elements and their target genes. Although data is increasingly being generated by us ^15;16^ and others ^29;35^, and has been incorporated into VAMPIRE where available, interrogations in a cell-type- or state-specific manner are still much needed. For instance, our recent work has demonstrated cell-type or tissue specific FIREs ^17;40^ and super interactive promoters (SIP)^41^ play key regulatory role and aid the identification and prioritization of functional regulatory elements and their corresponding genes. As more experimental data are generated, we will update VAMPIRE accordingly. Second, our list of 148,019 variants derives primarily from fine-mapping studies, which may be inaccurate in loci where more than one independent or partially independent signals exist. However, this limitation cannot be resolved before more powerful methods are developed for fine-mapping analysis for trans-ethnic GWAS. Finally, most of the annotations are based on analyses in European ancestry individuals (e.g. eQTL, pQTL, chromatin conformation etc.). Many ongoing efforts including ours are generating resources for non-European ancestry samples. For example, we are involved in several recently funded efforts to generate RNA-sequencing data in non-European ancestry individuals in hematopoietic cell types and anticipate relevant eQTL and sQTL annotations being added to VAMPIRE in upcoming years.

In conclusion, we have built a comprehensive annotation tool, VAMPIRE, which provides characterization and prioritization of blood cell trait related GWAS signals. Our results using existing functional experiments demonstrate that variants and genes prioritized by VAMPIRE are significantly more likely to be functionally validated at either the variant, gene, or variant-gene pair level. Annotation tools like VAMPIRE, which could be easily modified to apply to additional complex traits and diseases, are necessary to translate knowledge of GWAS significant variants to target genes and biological insights, and to guide our decisions to prioritize experimental validations of most likely functional regulatory variants/elements and their effector genes.

## Appendix

A1. Supplementary methods.

## Declaration of Interests

The authors declare no competing interests.

## Acknowledgement

This work was supported by the National Center for Advancing Translational Sciences, National Institutes of Health [R01HL146500 to A.P.R., R01HL129132 to Y.L., KL2TR002490 to L.M.R., R01DK103794 to V.G.S.], and the New York Stem Cell Foundation. C.A.C. and Y.L. are also partially supported by R01GM105785 and U01DA052713. V.G.S. is a New York Stem Cell Foundation-Robertson Investigator. The content is solely the responsibility of the authors and does not necessarily represent the official views of the NIH.

We would like to thank the Blood Cell Consortium (BCX) and the HemeNet investigators for comments on earlier versions of VAMPIRE. We want to thank Hanling Wang for visualization assistance. We also want to thank the Li lab members for feedback on the R Shiny app.

## Web Resources

VAMPIRE: http://shiny.bios.unc.edu/vampire/

GWAS summary statistics from Chen et al.^4^: http://www.mhi-humangenetics.org/en/resources/

GWAS Catalog: https://www.ebi.ac.uk/gwas/

PheWAS website: http://pheweb.sph.umich.edu

## Data Availability

The data underlying this article are available in the article and in its online supplementary material.

## Notes

### Competing Interest Statement

The authors have declared no competing interest.

