## Supplementary methods for "From GWAS Variant to Function: a Study of ~148,000 Variants for Blood Cell Traits"

Annotated Hematological Trait GWAS

The variants used in this application are from recent publications relating to hematologic traits. The majority of the variants (N=148,019; >99%) are from a recently performed trans-ethnic meta-analysis for a battery of hematological traits in up to 746,667 participants, including up to 184,535 non-European ancestry individuals (15,171 African ancestry, 9,368 Hispanic/Latino, 151,807 East-Asian, 8,189 South-Asian) [1]. The trans-ethnic meta-analysis identified 5,552 trait-variant associations at a significance threshold of p < 5x10^-9^. For 3,552 loci in which conditional analysis identified a single genome-wide significant variant in a European ancestry-specific analysis [2], fine mapping results were generated for each trans-ethnic and ancestry-specific dataset using an approximate Bayesian approach to create 95% credible sets identified as being significantly associated with seven red blood cell indices (HCT-Hematocrit; HGB-Hemoglobin Concentration; MCH-Mean Corpuscular Hemoglobin; MCHC- Mean Corpuscular Hemoglobin Concentration; MCV-Mean Corpuscular Volume; RBC-Red Blood Cell Count; RDW-Red Blood Cell Distribution Width), six white blood cell indices (BASO- Basophil Count; EOS- Eosinophil Count; LYM- Lymphocyte Count; MONO- Monocyte Count; NEU- Neutrophil Count; WBC-White Blood Cell Count), and two platelet related indices (MPV- Mean Platelet Volume; PLT- Platelet Count). In addition, two low frequency variants (<0.1%) that were identified in an analysis of TOPMed reference panel-imputed genotype data with hematological traits (hemoglobin (HGB), hematocrit (HCT), and white blood cell count (WBC)) in ~21,600 African-ancestry and ~21,700 Hispanic/Latino individuals were included [3].

Functional Annotation Details

Our predicted functional coding variant category is defined as follows: high confidence LoF variants are defined by VEP annotations; influential missense variants are defined to be those with MetaSVM score > 0 [4]; inframe indels and synonymous variants are considered influential if fathmm_XF_coding_score>0.5 [5]. For 1D annotations, we used tissues relevant to a given hematological trait grouping. For instance, we included ATAC-seq peak information, GenoSkyline+ scores, and key histone ChIP-seq peaks in spleen tissue, erythroid cells, and K562 cell lines for red blood cell related indices; fetal thymus tissue, GM12878 cells, macrophage cells, and T-cells and B-cells for white blood cell related indices; and megakaryocyte cells for platelet related indices.
